# A KDELR-mediated ER retrieval system guides the signaling pathways of UPR and AMPK to maintain cellular homeostasis

**DOI:** 10.1101/2023.06.14.545011

**Authors:** Mengdan Zhu, Zheng Fang, Yifan Wu, Fenfen Dong, Yuzhou Wang, Fan Zheng, Xiaopeng Ma, Shisong Ma, Jiajia He, Donald L. Hill, Xing Liu, Xuebiao Yao, Chuanhai Fu

## Abstract

KDELR (Erd2 in yeasts) mediates the retrieval of ER-resident proteins from the Golgi apparatus, yet how the KDELR-mediated ER retrieval system is involved in regulating cellular homeostasis has been unclear. Here we report that the loss of the Erd2-mediated ER retrieval system induces the unfolded protein response (UPR) and increases mitochondrial respiration and reactive oxygen species (ROS) in an UPR-dependent manner. Moreover, transcriptomic analysis revealed that expression of the genes involved in mitochondrial respiration and the tricarboxylic acid cycle is enhanced in an UPR-dependent manner in cells lacking Erd2. In cells lacking Erd2, the enhancement of mitochondrial respiration and ROS is required for maintaining cell viability. The loss of the Erd2-mediated ER retrieval system also activates AMPK, and consequently derepresses carbon catabolite repression. Hence, our work establishes a role of the KDELR/Erd2-mediated ER retrieval system in guiding the signalling pathways of AMPK and UPR and underscores the crucial role of the KDELR/Erd2-mediated ER retrieval system in the maintenance of cellular homeostasis.

## Introduction

ER-luminal proteins are required for maintaining ER homeostasis. A group of ER-luminal proteins, including the ER-stress response chaperones glucose-regulated protein-78 (GRP78) and glucose-regulated protein-94 (GRP94) and protein disulfide isomerase (PDI), carry the characteristic motif of xDEL (x, Lysine, Histidine, or Arginine; D, Aspartic Acid; E, Glutamic Acid; L, Leucine) at their C-terminal ends (Capitani and Sallese, 2009; Gerondopoulos et al., 2021). The characteristic motif enables the escaped ER-luminal proteins to be retrieved from the Golgi apparatus by the evolutionarily conserved receptor proteins, i.e., KDELR1/2/3 (KDEL receptors 1, 2, and 3) in humans and Erd2 (ER retention defective 2) in yeasts (Capitani and Sallese, 2009). Therefore, the KDELR/Erd2 retrieval system is involved in maintaining ER homeostasis. Although much progress has been made in delineating the mechanism of action of KDELR/Erd2, how the KDELR/Erd2-mediated retrieval system is involved in regulating cellular homeostasis has not been well understood.

Since its identification (Hardwick et al., 1990; Lewis and Pelham, 1990; Lewis et al., 1990; Semenza et al., 1990), KDELR/Erd2 has been characterized intensively (Aoe et al., 1997; Brauer et al., 2019; Cabrera et al., 2003; Cancino et al., 2014; Gerondopoulos et al., 2021; Lewis and Pelham, 1992; Scheel and Pelham, 1998; Semenza and Pelham, 1992; Townsley et al., 1994; Townsley et al., 1993; Trychta et al., 2018; Wilson et al., 1993). These efforts establish that KDELR/Erd2 is a seven-transmembrane transporter-like protein, cycling between the Golgi apparatus and the ER to retrieve the xDEL-containing ER resident proteins. In addition, it has been demonstrated that on the surface of KDELR/Erd2, a polar luminal cavity is present, responsible for the recognition and selection of the xDEL-containing ER resident proteins (Brauer et al., 2019; Gerondopoulos et al., 2021).

*Erd2* in the budding yeast *Saccharomyces cerevisiae* is essential, and Erd2-deficient cells accumulate intercellular membranes and lipid droplets and have impaired secretory pathways (Semenza et al., 1990). Despite the essentiality of *ERD2*, the Erd2-mediated retrieval system does not appear to be essential for cell viability (Hardwick et al., 1992; Semenza et al., 1990). These paradoxical findings are interpreted to be due to the existence of the Ire1-mediated mechanism that works in parallel with the Erd2-retrieval system to maintain ER homeostasis (Beh and Rose, 1995). Ire1 (Inositol-requiring enzyme 1) is an ER transmembrane sensor responsible for activating the unfolded protein response (UPR) signaling pathway (Wu et al., 2014). Given the role of Erd2 in ER quality control, impairment of Erd2 may also induce UPR. This hypothesis has not been tested for yeasts. Nonetheless, impairment of the KDELR/Erd2 retrieval system in mammalian cells enhances UPR induced chemically (Wu et al., 2014). The cellular responses caused by the impairment of KDELR/Erd2 and the related physiological significance await further investigation.

Impairment of the KDELR/Erd2 retrieval system has pathophysiological consequences. KDELR1 mutations incapable of binding ligands reduce the number of naive T cells (Kamimura et al., 2015) or cause lymphopenia (Siggs et al., 2015). In addition, mice carrying the KDELR1 mutant D193N, capable of ligand recognition but defective in distribution to the ER, display accumulation of misfolded proteins in the ER and develop dilated cardiomyopathy (Hamada et al., 2004). These pathophysiological consequences appear to be due to the responses of stress signaling, including the integrated stress responses (ISR) and/or the UPR (Hamada et al., 2004; Kamimura et al., 2015; Yamamoto et al., 2003), implying that a profound metabolic remodeling occurs upon impairment of the KDELR/Erd2 retrieval system. How the KDELR/Erd2 retrieval system mobilizes organelles, including the ER and mitochondria, to remodel metabolism remains unclear.

Using the fission yeast *Schizosaccharomyces pombe* as a model organism (Chiron et al., 2007; Dong et al., 2022; Zheng et al., 2019), we performed a microscopy-based screen and has identified Emr1 as a crucial protein regulating the contact between mitochondria and the ER (Rasul et al., 2021). In the screen, we also identified the KDELR counterpart Erd2 as a protein regulating mitochondrial function. In fission yeast, Erd2 has not been well characterized. However, it has been shown that *erd2* is not essential and cells lacking Erd2 are sensitive to hydrogen peroxide, caffeine, and tunicamycin (Calvo et al., 2009; Guydosh et al., 2017). In the present work, we show that the absence of Erd2 induces UPR and increases mitochondrial respiration and ROS in an UPR-dependent manner. The absence of Erd2 affects mitochondrial functions by increasing expression of the genes involved in mitochondrial respiration and the tricarboxylic acid cycle (TCA), and increases in mitochondrial respiration and ROS are required for maintaining cell viability. We further demonstrate that the absence of Erd2 activates AMPK and derepresses carbon catabolite repression. Hence, the present work reveals an uncharacterized role of the KDELR/Erd2 retrieval system in maintaining cellular homeostasis by guiding the signaling pathways of UPR and AMPK.

## Results

### Erd2 is required for maintaining mitochondrial tubular structures and for regulating mitochondrial respiration and ROS

Erd2 resides mainly at the Golgi apparatus and cycles between the Golgi apparatus and the ER (Robinson and Aniento, 2020). The canonical role of Erd2 is the retrieval of the ER resident proteins that escape from the ER (Capitani and Sallese, 2009). Intriguingly, we found that the absence of *erd2* caused mitochondrial fragmentation (Fig. 1A). Thus, it is puzzling how a protein that does not appear to localize to mitochondria is involved in regulating mitochondrial functions.

**Fig. 1.**
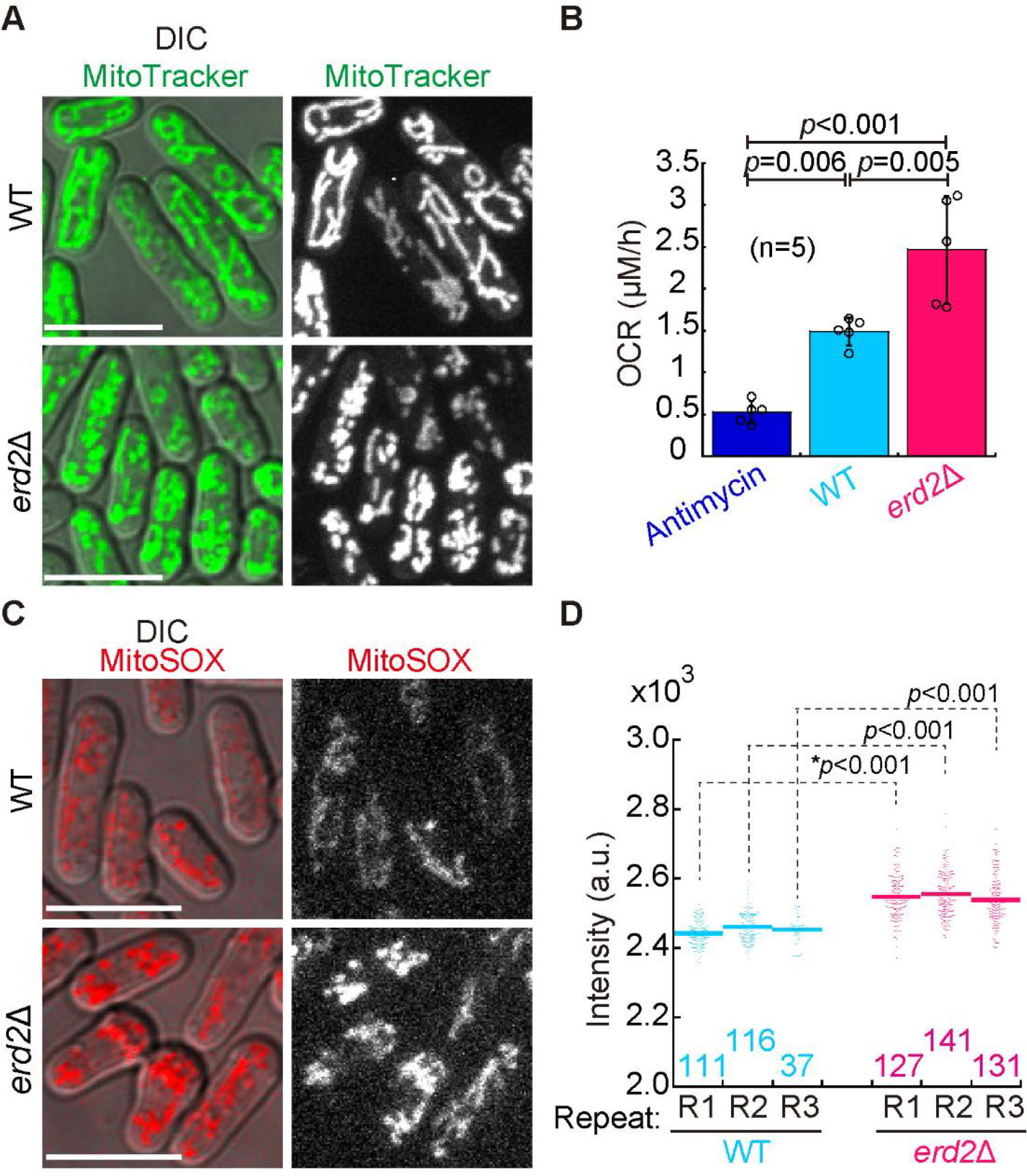
The absence of Erd2 caused mitochondrial fragmentation and increased mitochondrial respiration and ROS. (A) Maximum projection images of WT and *erd2Δ* cells cultured in EMM media. Mitochondria were stained with MitoTracker Green. Note that mitochondria became fragmented in the absence of Erd2. *DIC*, differential interference contrast. Scale bar, 10 μm. (B) Oxygen consumption rates (OCR) of the indicated cells. A Strathkelvin oxygen respirometer was used to measure OCR. Shown is a representative result from three independent experiments (also see Fig. S1 A for all data), and 5 repeats (*n*) were performed for each type of strains. The height of the column is the mean, and error bars indicate S.D. The *p* values were calculated by one-way ANOVA with a post hoc Tukey HSD test. Antimycin A (0.15 µg/ml) was used to inhibit mitochondrial respiration. Note that the absence of Erd2 increased OCR. (C) Assessment of mitochondrial reactive oxygen species (ROS) of WT and *erd2Δ* cells by staining with MitoSOX Red. *DIC*, differential interference contrast. Scale bar, 10 μm. (D) Quantification of average fluorescent intensity of mitochondrial staining indicated in (C). Three independent experiments were performed (indicated by R1-3), and the number of cells analyzed is indicated. The *p* values were calculated by the Wilcoxon-Mann-Whitney rank sum test or Student’s *t*-test (indicated by asterisks). Note that the absence of Erd2 increased ROS.

To address the above question, we began by examining mitochondrial morphology and functions in the prototrophic wild-type (WT) and *erd2*-deleted (*erd2*Δ) strains. Staining the prototrophic strains with MitoTracker Green showed that mitochondria became highly fragmented in *erd2*Δ cells cultured in the Edinburgh Minimal Medium (EMM) (Fig. 1 A). Moreover, the oxygen consumption rate of *erd2*Δ cells was higher than that of WT cells (Fig. 1 B and Fig. S1 A), suggesting that, in *erd2*Δ cells, mitochondrial respiration was increased. In addition, MitoSOX staining was employed to assess reactive oxygen species (ROS) within mitochondria. As shown in Fig. 1 C and D, mitochondrial ROS was higher in *erd2*Δ cells than in WT cells. Collectively, these results suggest that Erd2 is involved in regulating mitochondrial morphology, mitochondrial respiration, and mitochondrial ROS.

### Erd2 guides the signaling pathway of UPR

Given the canonical role of Erd2 in maintaining ER homeostasis, we reasoned that the absence of Erd2 may regulate mitochondria via the ER. To test this hypothesis, we assessed the unfolded protein response (UPR) signaling pathway. Since reduced expression of *gas2* and *yop1* is a hallmark of UPR in fission yeast (Kimmig et al., 2012), we monitored the expression of *gas2* and *yop1* by quantitative reverse transcription PCR (RT-qPCR). As shown in Fig. 2 A, similar to the effect by treatment of cells with the UPR inducer tunicamycin (0.5 µg/ml for 1 hour), the absence of Erd2, but not the absence of the UPR activator Ire1, reduced the expression of both *gas2* and *yop1*. The absence of both Erd2 and Ire1, however, did not reduce the expression of *gas2* and *yop1*. These results suggest that the absence of Erd2 induces UPR. Cells defective in UPR are sensitive to tunicamycin (Kimmig et al., 2012). Consistently, *ire1*Δ cells failed to grow on EMM plates containing tunicamycin (Fig. 2 B). Paradoxically, *erd2*Δ cells, in which UPR was activated (Fig. 2 A), also grew poorly on EMM plates containing tunicamycin. Considering the canonical role of Erd2 in retrieving the ER-resident proteins required for maintaining ER homeostasis (Capitani and Sallese, 2009), we favored the interpretation that the poor cell growth was due to impairment of the ER protein-folding capacity caused by the failure in the retrieval of ER-resident proteins in *erd2*Δ cells. Under unstressed conditions, i.e., when cells were grown on EMM plates, *ire1*Δ and *erd2*Δ had no noticeable effect on cell growth but *ire1*Δ*erd2*Δ impaired cell growth (Fig. 2 B). Collectively, these results indicate that when the KDELR/Erd2 retrieval system is impaired under unstressed conditions, UPR becomes necessary for maintaining cellular homeostasis.

**Fig. 2.**
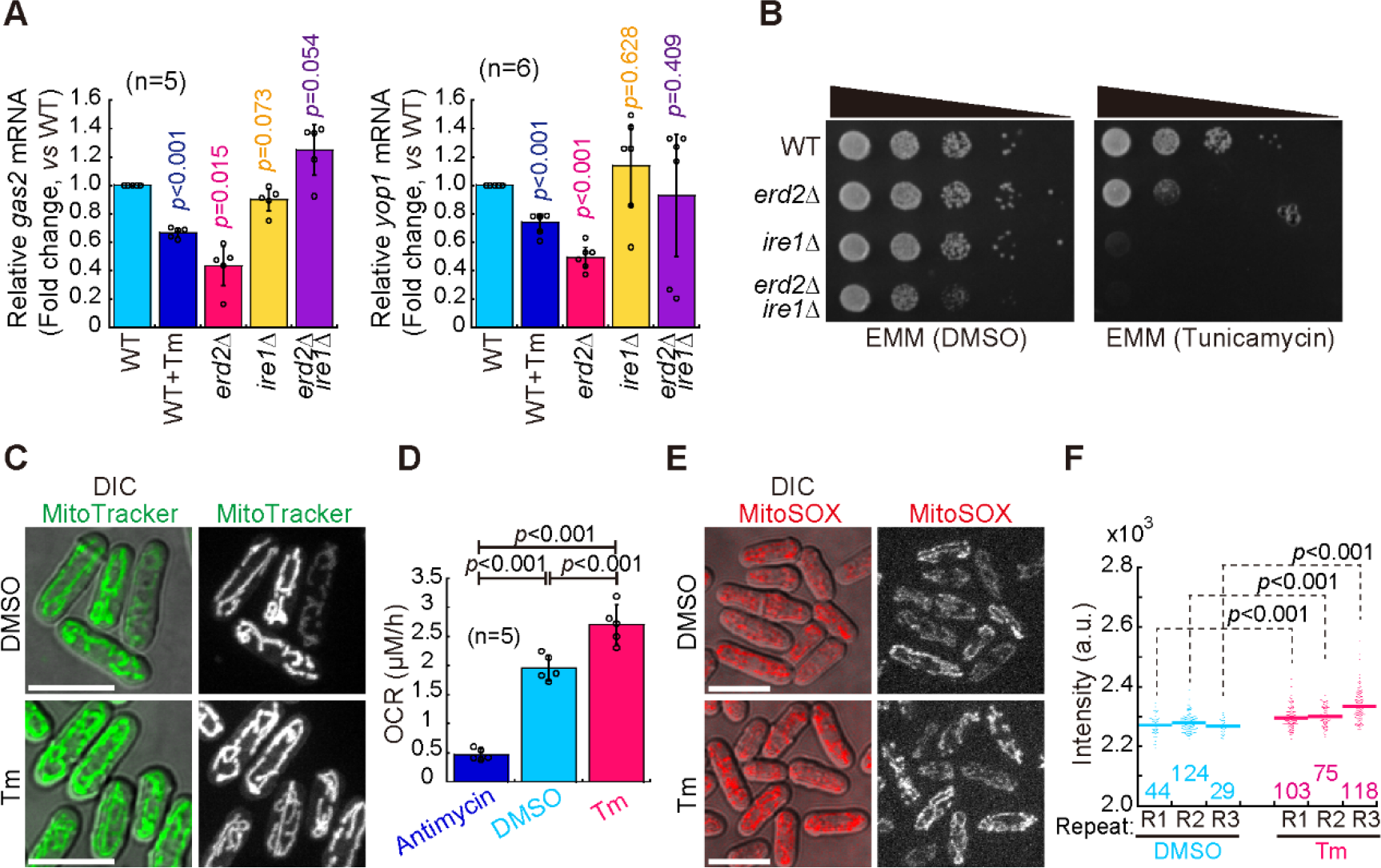
The absence of Erd2 activated UPR, and UPR increased mitochondrial respiration and ROS. (A) The expression of *gas2* and *yop1* in the indicated cells analyzed by real time quantitative PCR (RT-qPCR). Shown are fold changes of the indicated cells over wild-type control. The number (*n*) of the independent measurements is indicated. The height of each bar represents the mean, and error bars indicate confidence intervals (alpha=0.05). Single-group Student’s *t*-test was used to calculate the *p* values. (B) Cell growth assays. The indicated cells, by 10-fold serial dilutions, were spotted on EMM plates containing tunicamycin or DMSO. Images were taken after three days of culture at 30 ℃. (C) Maximum projection images of WT cells treated with DMSO (solvent) or tunicamycin (0.5 μg/ml) for 1 hour. Mitochondria were stained with MitoTracker Green. Note that, in cells treated with tunicamycin (Tm), mitochondria remained tubular. *DIC*, differential interference contrast. Scale bar, 10 μm. (D) Oxygen consumption rates of the indicated cells. Shown is a representative result from three independent experiments (also see Fig. S1 B for all data), and 5 repeats were performed for each type of treatment. The height of the column is the mean, and error bars indicate S.D. The *p* values were calculated by one-way ANOVA with a post hoc Tukey HSD test. Antimycin A was used to inhibit mitochondrial respiration. Note that tunicamycin increased OCR. (E) Assessment of mitochondrial ROS of DMSO- or tunicamycin-treated cells by staining with MitoSOX Red. *DIC*, differential interference contrast. Scale bar, 10 μm. (F) Quantification of average fluorescent intensity of mitochondrial staining indicated in (E). Three independent experiments were performed (indicated by R1-3), and the number of cells analyzed is indicated. The *p* values were calculated by a Wilcoxon-Mann-Whitney rank sum test. Note that tunicamycin increased ROS.

### UPR increases mitochondrial respiration and mitochondrial ROS

Next, we tested whether UPR affects mitochondria. To induce UPR, we treated cells with 0.5 µg/ml tunicamycin. MitoTracker Green staining showed that tunicamycin did not alter mitochondrial tubular structures during a 1-hour treatment with tunicamycin (Fig. 2 C). By contrast, tunicamycin increased the oxygen consumption rate of cells (Fig. 2 D and Fig. S1 B), which was similar to the effect caused by the absence of Erd2 (Fig. 1 B). Moreover, tunicamycin increased ROS within mitochondria (Fig. 2 E and F), which was similar to the effect caused by the absence of Erd2 (Fig. 1 C and D). Hence, these results suggest that UPR increases mitochondrial respiration and ROS but does not affect mitochondrial morphology.

### Erd2 regulates mitochondrial respiration and ROS production via UPR

To test whether the mitochondrial alteration caused by the absence of Erd2 depends on UPR, we compared mitochondrial morphology, mitochondrial respiration, and mitochondrial ROS in WT, *erd2*Δ, *ire1*Δ, and *erd2*Δ*ire1*Δ cells. As shown in Fig. 3 A, mitochondria were fragmented in *erd2*Δ cells, but not in WT, *ire1*Δ, or *erd2*Δ*ire1*Δ cells. This result suggests that the absence of Erd2 caused mitochondrial fragmentation in an Ire1-dependent manner. Of note, the absence of Erd2 resulted in UPR activation (Fig. 2 A), but induction of UPR by tunicamycin did not cause mitochondrial fragmentation (Fig. 2 C). Therefore, mitochondrial fragmentation caused by the absence of Erd2 requires UPR, but UPR alone is not sufficient to induce mitochondrial fragmentation.

**Fig. 3.**
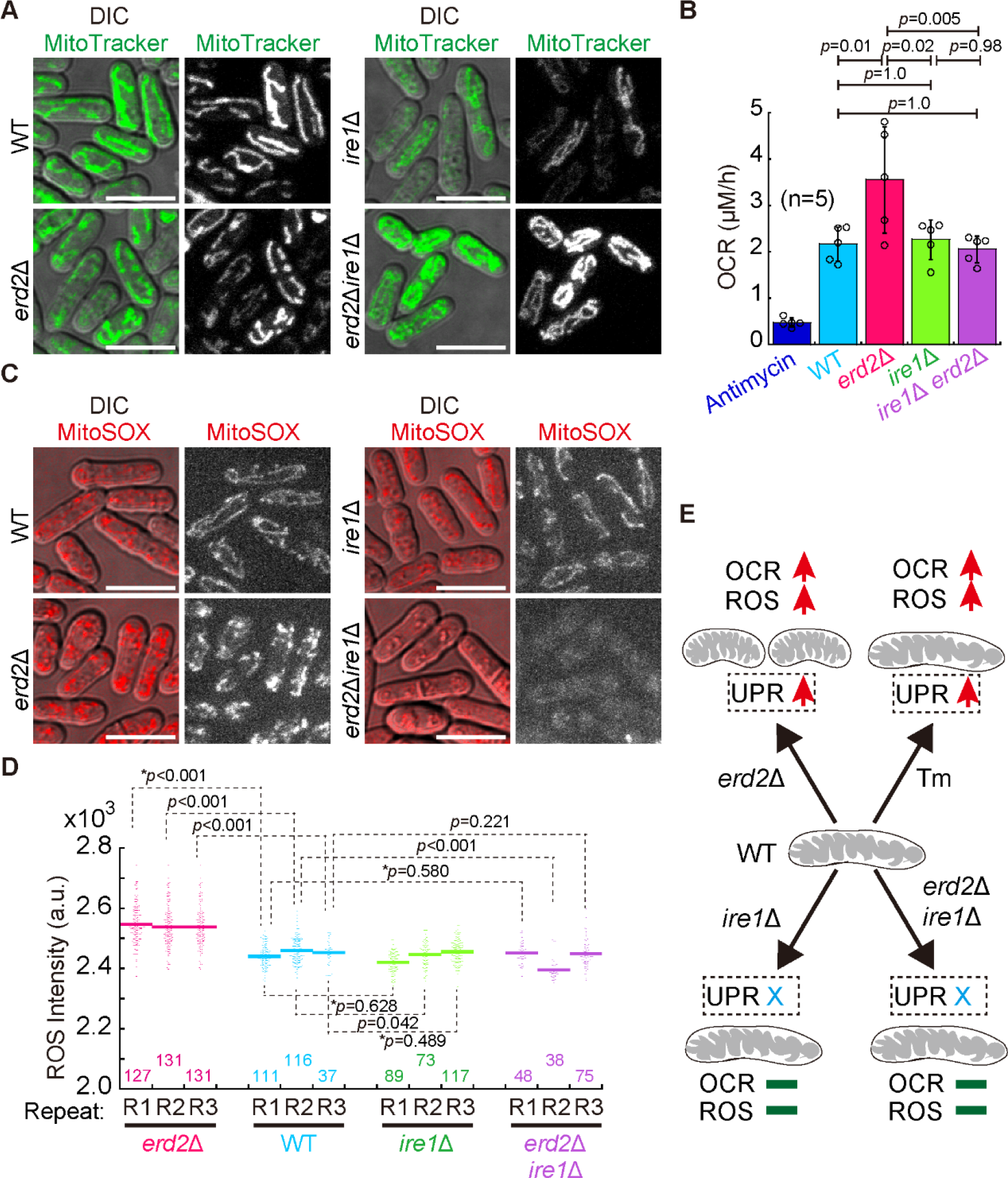
Increases in mitochondrial respiration and ROS caused by the absence of Erd2 depended on Ire1. (A) Maximum projection images of WT, *erd2Δ*, *ire1Δ*, and *erd2Δire1Δ* cells stained with MitoTracker Green. Note that mitochondria were fragmented in *erd2Δ* cells and became tubular in *erd2Δire1Δ* cells. *DIC*, differential interference contrast. Scale bar, 10 μm. (B) Oxygen consumption rates (OCR) of the indicated cells. Shown is a representative result from three independent experiments (also see Fig. S3 C for all data), and 5 repeats were performed for each type of strain. The height of the column is the mean, and error bars indicate S.D. The *p* values were calculated by one-way ANOVA with a post hoc Tukey HSD test. Antimycin A was used to inhibit mitochondrial respiration. Note that the absence of Erd2, but not the absence of both Erd2 and Ire1, increased OCR. (C) Assessment of mitochondrial ROS of the indicated cells by staining with MitoSOX Red. *DIC*, differential interference contrast. Scale bar, 10 μm. (D) Quantification of average fluorescent intensity of mitochondrial staining indicated in (C). Three independent experiments were performed (indicated by R1-3), and the number of cells analyzed is indicated. The *p* values were calculated by a Wilcoxon-Mann- Whitney rank sum test or by Student’s t-test (indicated by asterisks). Note that the absence of Erd2, but not the absence of both Erd2 and Ire1, increased ROS. (E) Summary of the effects of *erd2Δ*, *ire1Δ*, *ire1Δerd2Δ*, and tunicamycin on mitochondrial morphology, respiration, and ROS. Red arrows indicate an increase, blue *X* indicates no response of UPR, and green bars indicate no significant change.

We further measured the oxygen consumption rates of WT, *erd2*Δ, *ire1*Δ, and *erd2*Δ*ire1*Δ cells (Fig. 3 B and Fig. S1 C). The results showed that the oxygen consumption rate of *erd2*Δ cells, but not *ire1*Δ or *erd2*Δ*ire1*Δ cells, was increased. Of note, induction of UPR by tunicamycin also increased the oxygen consumption rate (Fig. 2 D). Therefore, the absence of Erd2 increased mitochondrial respiration in an UPR/Ire1-dependent manner. Similarly, the absence of Erd2 increased mitochondrial ROS in an UPR/Ire1-dependent manner (Fig. 3 C and D). The alterations of mitochondria by these mutants and tunicamycin are summarized in Fig. 3 E. Collectively, we concluded that the absence of Erd2 increases mitochondrial respiration and ROS in a UPR-dependent manner.

### Erd2 regulates expression of the genes involved in the mitochondrial electron transport chain and the tricarboxylic acid cycle

To reveal how the absence of Erd2 increases mitochondrial respiration and mitochondrial ROS, we performed RNA-sequencing for WT, *erd2*Δ, *ire1*Δ, and *erd2*Δ*ire1*Δ cells (samples in triplicate). GO (gene ontology) analysis of the differentially expressed genes (*erd2*Δ vs WT) revealed that genes related to the mitochondrial electron transport chain and the tricarboxylic acid cycle were enriched (Fig. S2). Examination of the GO-enriched genes revealed that the absence of Erd2 increased (with *P* values (adjusted) <0.05 and fold change >1.5) the expression of 1, 2, 8, 6, 13 genes encoding proteins in the yeast counterpart of the NADH dehydrogenase (complex I), succinate dehydrogenase (complex II), oxidoreductase (complex III), cytochrome c oxidase (complex IV), and the ATP synthase (complex V), respectively (Fig. 4 A). Therefore, expression of ∼80% of the genes in the mitochondrial respiratory chain was increased, consistent with the enhancement of mitochondrial respiration in *erd2*Δ cells (Fig. 3 B). Moreover, expression of most of the genes encoding the proteins responsible for the tricarboxylic acid cycle were also increased (Fig. 4 A). Such increased expression of the genes regulating the mitochondrial respiratory chain and the tricarboxylic acid cycle was not found in *ire1*Δ (Fig. S3) and *erd2*Δ*ire1*Δ cells (Fig. S4). RT-qPCR was conducted to assess the expression of the representative genes *atp1* (complex V), *atp2* (complex V), *cox13* (complex IV), and *rip1* (complex III) in WT, *erd2*Δ, *ire1*Δ, and *erd2*Δ*ire1*Δ cells, confirming the above RNA-sequencing findings (Fig. 4 B). These results suggest that the absence of Erd2 increases expression of the genes regulating mitochondrial respiration and the tricarboxylic acid cycle in a UPR/Ire1-dependent manner.

**Fig. 4.**
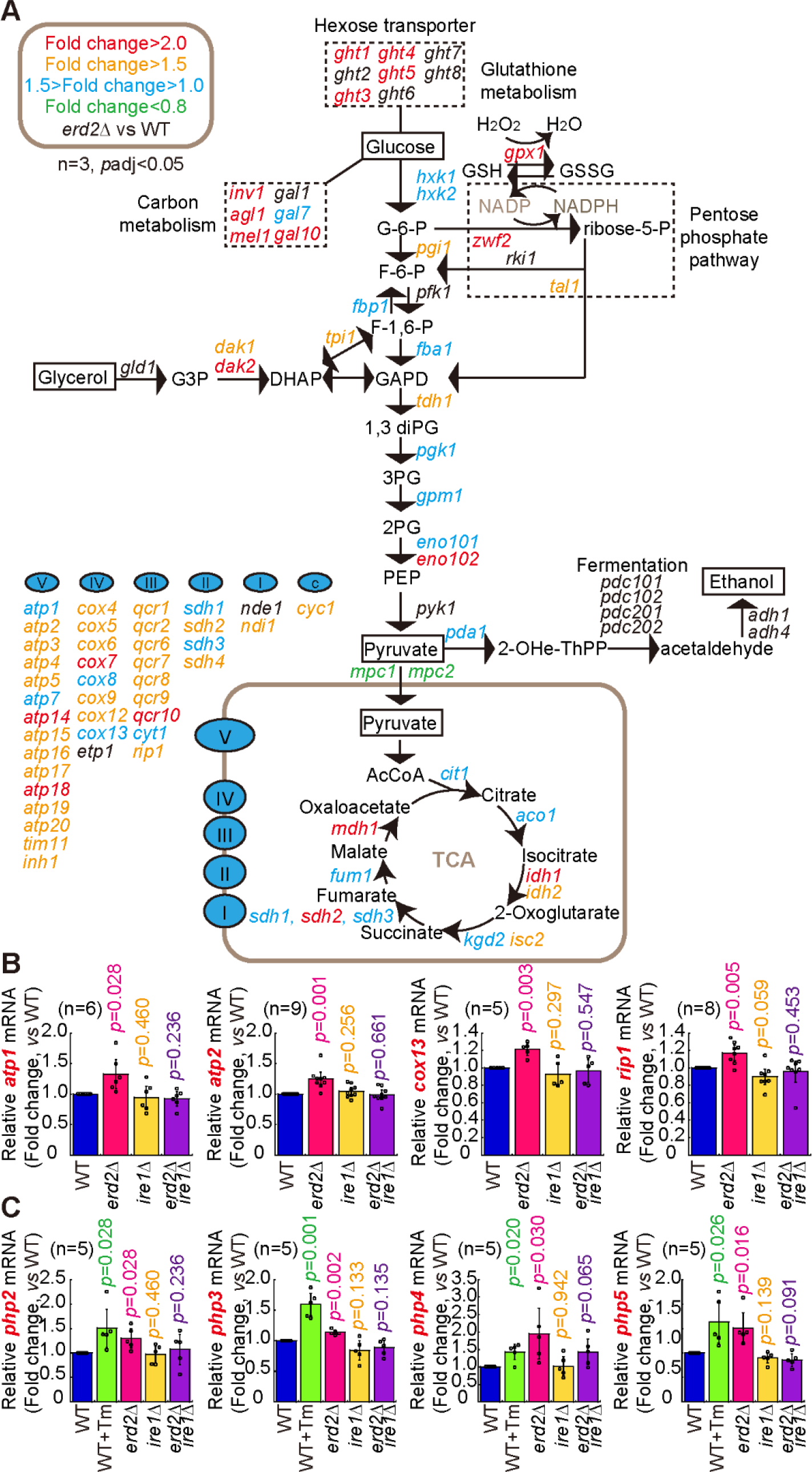
The absence of Erd2 increased expression of the genes involved in mitochondrial respiration and the tricarboxylic acid cycle in an *ire1*-dependent manner. (A) Diagram illustrating glycolysis, the mitochondrial electron transport chain (ETC), and the tricarboxylic acid cycle (TCA). Indicated, in red, orange, blue, and green colors, are the genes that exhibit differential expression in *erd2Δ* cells (vs WT, adjusted *p*-values (padj)<0.05). The genes of the mitochondrial electron transport chain and the ATP synthetase are listed. For the comparisons between *erd2Δ* and *erd2Δire1Δ* and WT, see supplementary Fig. S3 and Fig S4. (B) RT-qPCR was conducted to analyze the expression of the representative ETC genes *atp1*, *atp2*, *cox13*, and *rip1* in the indicated cells. Shown are fold changes of the tested genes in the indicated cells (vs WT). The number (*n*) of experimental replicates is indicated. The height of each bar represents the mean, and error bars indicate confidence intervals (alpha=0.05). A single-group Student’s *t*-test was used to calculate the *p* values. (C) RT-qPCR was conducted to analyze the expression of the genes encoding the CCAAT-binding transcription factors (i.e. *php2, php3*, *php4,* and *php5*) in the indicated cells. Shown are fold changes of the tested genes in the indicated cells over a wild-type control. The number (*n*) of experimental replicates is indicated. The height of each bar represents the mean, and error bars indicate confidence intervals (alpha=0.05). A single-group Student’s *t*-test was used to calculate the *p* values. *Tm* indicates cells treated with tunicamycin to induce UPR.

In the budding yeast *Saccharomyces cerevisiae*, the HAP transcriptional complex, which binds to the CCAAT-binding site, promotes the expression of genes involved in the mitochondrial respiratory chain and the tricarboxylic acid cycle (Buschlen et al., 2003). Similarly, in the fission yeast *Schizosaccharomyces pombe*, the HAP component Php2 is involved in regulating expression of the subunit in the mitochondrial respiratory chain (Takuma et al., 2013). Therefore, we tested, by RT-qPCR, whether the absence of Erd2 affects the expression of the HAP complex, i.e., Php2-Php3-Php4-Php5, in fission yeast. As shown in Fig. 4 C, the absence of Erd2 increased the expression of *php2*, *php3*, *php4*, and *php5*. Induction of UPR by treatment of cells with (0.5 µg/ml) tunicamycin also increased the expression of *php2*, *php3*, *php4*, and *php5* (Fig. 4 C). Moreover, increased expression of *php2*, *php3*, *php4*, and *php5* was not found in *ire1*Δ or *erd2*Δ*ire1*Δ cells (Fig. 4 C). Collectively, these results suggest that the absence of Erd2 increases expression of the genes encoding the CCAAT-binding transcription complex in an UPR/Ire1-dependent manner.

Given the facts that the absence of Erd2 induces UPR (Fig. 2) and that the absence of Erd2 increases mitochondrial respiration and ROS in an UPR/Ire1-dependent manner (Fig. 3), we proposed the model presented in Fig. 8. The absence of Erd2 induces UPR, and consequently, increases the expression of genes encoding the CCAAT-binding transitional complex. The increased expression of CCAAT-binding transitional complex then promotes expression of the genes involved in the mitochondrial respiratory chain and the tricarboxylic acid cycle. As a result, mitochondrial respiration and ROS are enhanced in cells lacking Erd2.

### The absence of Erd2 activates AMPK

Expression of *ght5*, the hexose transporter gene, is repressed by the transcription factor Scr1 under glucose-rich conditions, and the repression effect is derepressed by AMPK. Upon glucose starvation, AMPK is activated and phosphorylates Scr1, causing the export of Scr1 from the nucleus and promoting the expression of *ght5*. Intriguingly, in our RNA-seq analysis (Fig. 4 A), we found that expression of genes encoding hexose transporters (i.e., *ght1*, *ght3*, *ght4*, and *ght5*) were significantly increased. This result prompted us to test whether the signaling pathway of AMPK is activated in cells lacking Erd2. We began by examining the localization of Scr1-GFP in WT and *erd2*Δ cells cultured in glucose-rich medium. Interestingly, Scr1-GFP localized mainly within the nucleus in WT cells but in the cytoplasm in *erd2*Δ cells (Fig. 5 A and B). The cytoplasmic localization of Scr1-GFP in *erd2*Δ cells was similar to that in WT cells that were cultured in glucose-poor medium (Fig. 5 A and B). We further assessed AMPK phosphorylation, an event indicative of AMPK activation, by western blotting analysis with an antibody used previously (Ling et al., 2020). As shown in Fig. 5 C and D, the phosphorylation levels of AMPK of *erd2*Δ cells were significantly higher than those of WT cells cultured in glucose-rich medium but lower than those of WT cells cultured in glucose-poor medium, indicative of partial activation of AMPK in cells lacking Erd2. Collectively, these results suggest that Erd2 guides the activation of AMPK.

**Fig. 5.**
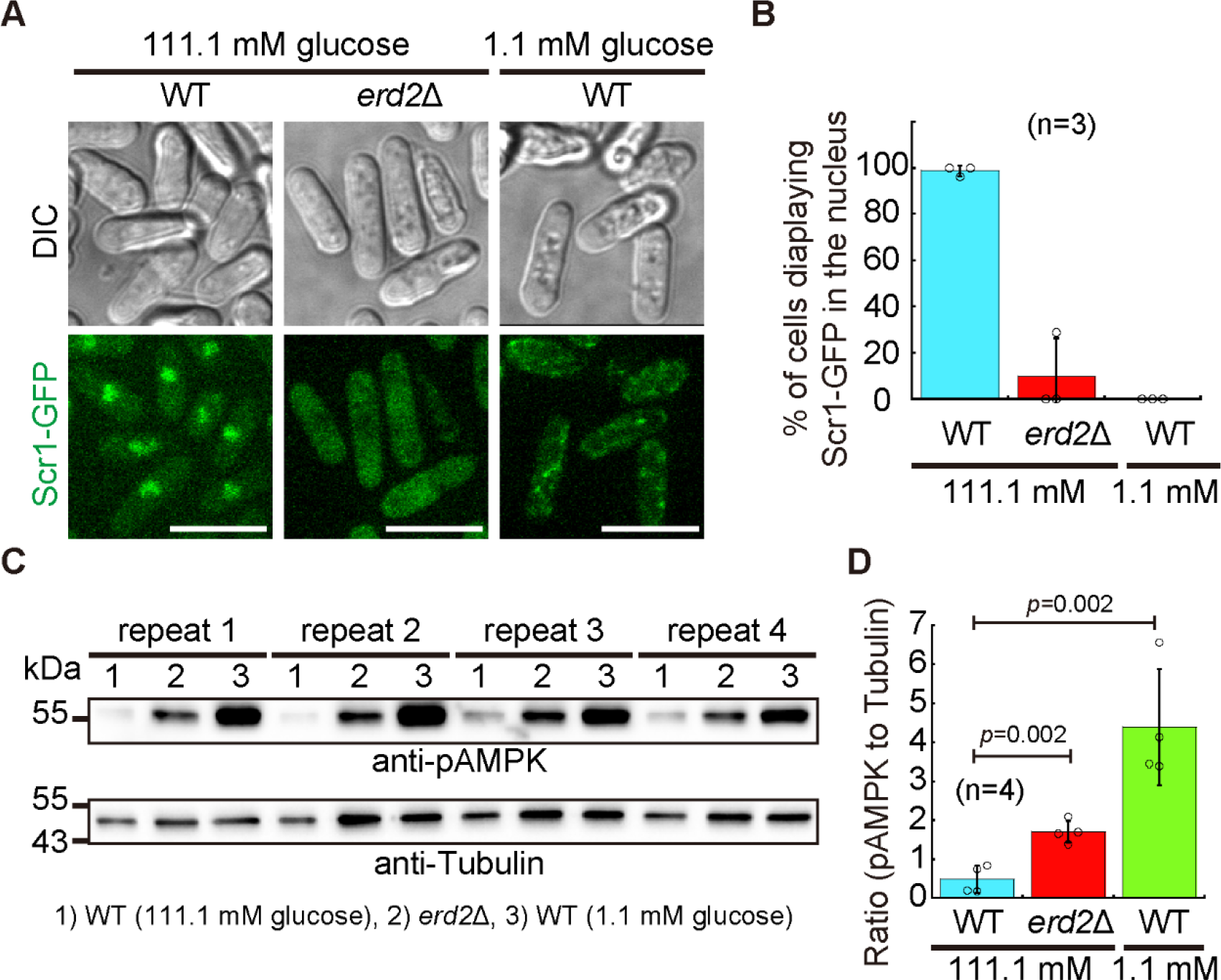
The absence of Erd2 caused delocalization of Scr1 and AMPK phosphorylation. (A) Maximum projection images of cells expressing Scr1-GFP. WT and *erd2Δ* cells cultured in glucose-rich (111.1 mM) EMM medium and WT cells cultured in glucose-limiting (1.1 mM) EMM medium were examined by spinning-disk microscopy. (B) Quantification of percentage of cells exhibiting Scr1-GFP in the nucleus. Three independent experiments were performed. (C) Western blotting analysis was performed to analyze the phosphorylation of AMPK for the three types of samples as indicated. Four repeats of the experiments are shown. (D) Quantification of the intensity ratio of pAMPK to Tubulin. The top of the column indicates the mean, and statistical analysis was performed by Student’s *t*-test.

### Increases in mitochondrial respiration and ROS are required to maintain viability of cells lacking Erd2

To understand the physiological significance of the increased mitochondrial respiration and ROS in cells lacking Erd2, we assessed cell viability by testing colony formation (Fig. 6 A). Specifically, cells at the exponential phase were treated with 1 mM Trolox (an antioxidant used to clean ROS), 0.15 µg/ml antimycin (an inhibitor of the mitochondrial respiratory chain), or the solvent DMSO for 4 or 8 hours, followed by single-cell dissection on EMM plates with a tetrad-dissection microscope (Fig. 6 A). Colony formation on the EMM plates was examined after three days. As shown in Fig. 6 B and D, treatment of WT cells with Trolox did not affect cell viability, whereas treatment of *erd2*Δ cells with Trolox, for either 4 or 8 hours, impaired cell viability. Similar results were found when cells were treated with antimycin for either 4 or 8 hours (Fig. 6 C and E). Thus, in cells lacking Erd2, the increases in mitochondrial respiration and ROS are required for maintaining cell viability.

**Fig. 6.**
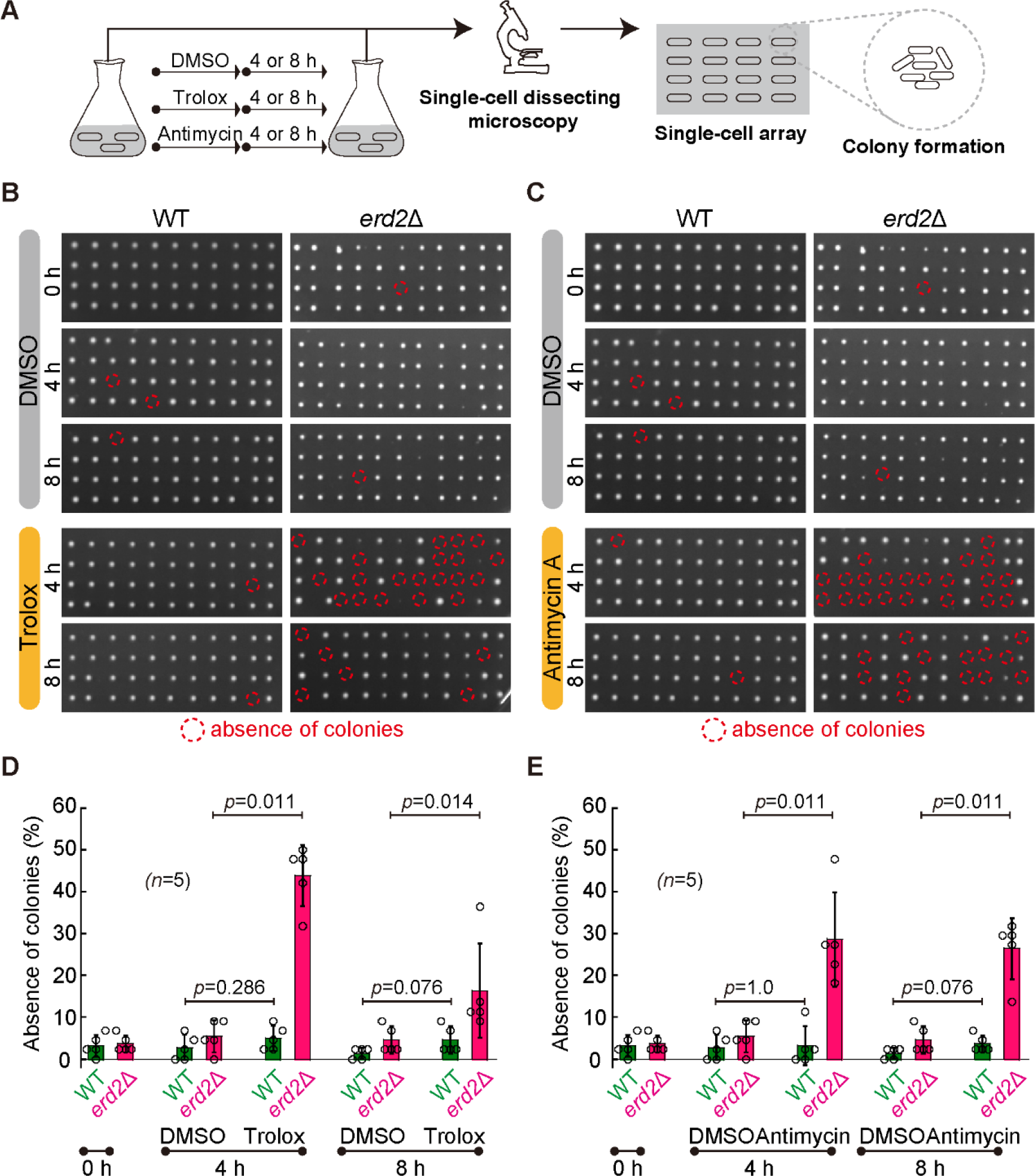
Increases in mitochondrial respiration and mitochondrial ROS production were required for maintaining viability of *erd2Δ* cells. (A) Diagram illustrating the procedure for assaying cell viability by tetrad-dissection analysis. The capability of forming a colony from a single cell after treatment with the indicated chemicals was examined on EMM plates. (B and C) Single-cell colony arrays. WT and *erd2Δ* cells were treated with (1 mM) Trolox (B), (0.15 µg/ml) antimycin (D), or DMSO (solvent) for 4 or 8 hours. Cells without treatment are controls. Dashed circles indicated the absence of colonies, i.e., cell death. (D and E) Quantification of the percentage of cell death (the absence of colonies). The top of the column is the mean, and error bars indicate S.D. Circles indicate the value measured from each experiment, and five independent experiments were performed. Student’s *t*-test was used to calculate the *p* values.

### AMPK activation is required to maintain viability of cells lacking Erd2

To understand the physiological significance of AMPK activation in cells lacking Erd2, we similarly assessed cell viability by testing colony formation. We began by testing the potent inhibitor of AMPK, i.e., BAY3827 (Lemos et al., 2021). Treatment of *erd2*Δ cells with 10 µM BAY3827 for 2 hours was enough to inhibit the export of Scr1-GFP (Fig. 7 A and B), suggesting that BAY3827 was functional in inhibiting AMPK in fission yeast. Colony formation assays exhibited that treatment of WT cells with BAY3827 did not affect cell viability, whereas treatment of *erd2*Δ cells with BAY3827, for either 4 or 8 hours, significantly impaired cell viability (Fig. 7 C and D). Moreover, deletion of both *erd2* and *ssp2* (the catalytic component of the AMPK complex) exhibited an additive effect on the impairment of cell growth (Fig. 7 E). Collectively, we concluded that in cells lacking Erd2, AMPK activation is required for maintaining cell viability.

**Fig. 7.**
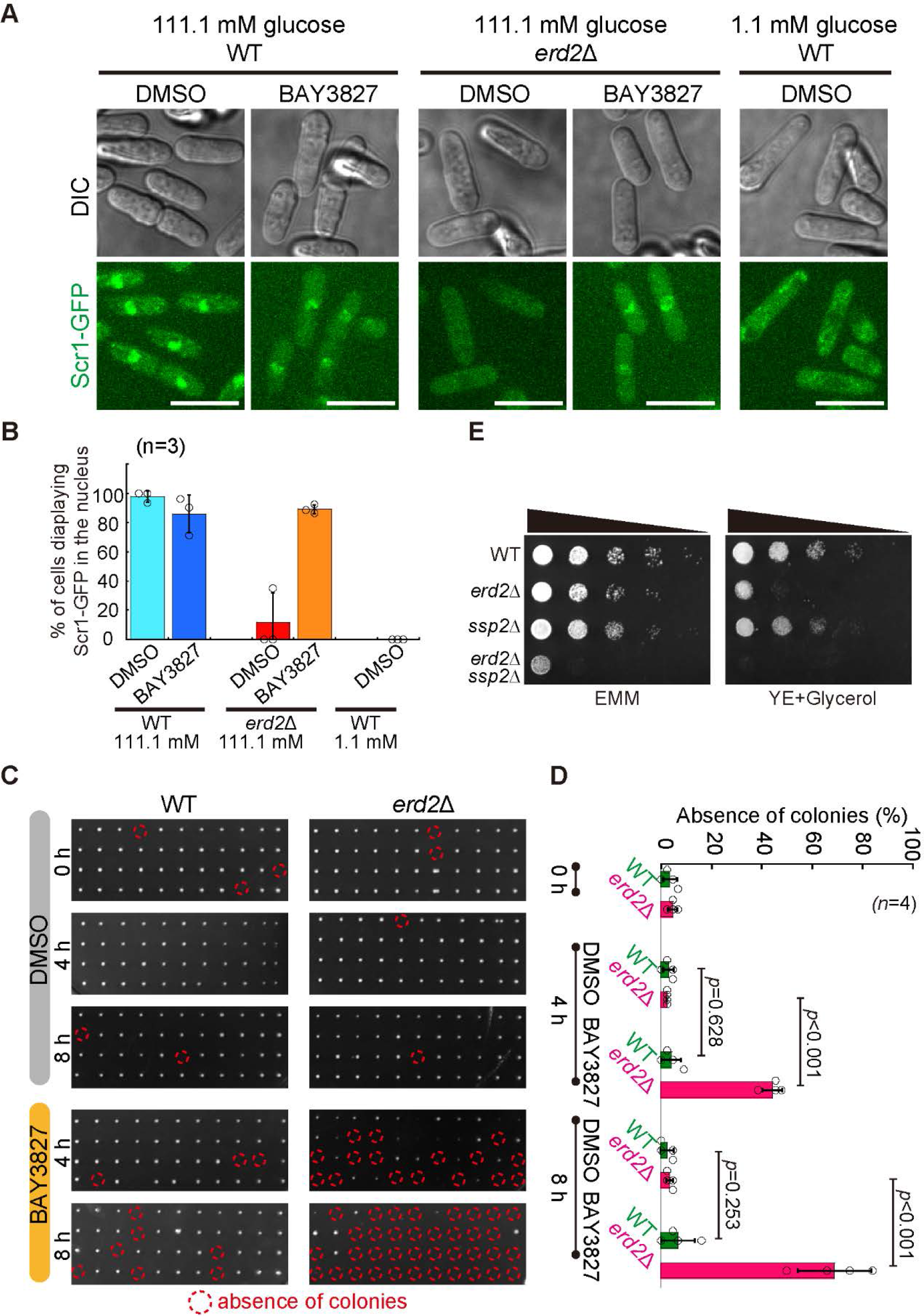
AMPK activation was required for maintaining viability of *erd2Δ* cells. (A) Maximum projection images of cells expressing Scr1-GFP. WT and *erd2Δ* cells cultured in glucose-rich (111.1 mM) EMM medium and WT cells cultured in glucose-limiting (1.1 mM) EMM medium were examined by spinning-disk microscopy. BAY38827 at 10 µM or DMSO was used to treated cells for 2 hours. (B) Quantification of percentage of cells exhibiting Scr1-GFP in the nucleus. Three independent experiments were performed. (C) Single-cell colony arrays. WT and *erd2Δ* cells were treated with (10 µM) BAY3827 for 4 or 8 hours. Cells without treatment are controls. Dashed circles indicated the absence of colonies, i.e., cell death. (D) Quantification of the percentage of cell death (the absence of colonies). The top of the column is the mean, and error bars indicate S.D. Circles indicate the value measured from each experiment, and four independent experiments were performed. Student’s *t*-test was used to calculate the *p* values. (E) Cell growth assays. The indicated cells, by 10-fold serial dilutions, were spotted on EMM plates or nonfermentable YE plates containing glycerol. Images were taken after three days of culture at 30 ℃.

## Discussion

The KDELR/Erd2 retrieval system is involved in maintaining protein homeostasis in the ER by retrieving the xDEL-containing ER resident proteins that escape from the ER (Brauer et al., 2019; Capitani and Sallese, 2009). In the present work, we report that Erd2 plays a crucial role in maintaining cellular homeostasis by guiding the signaling pathways of UPR and AMPK (Fig. 8).

**Fig. 8.**
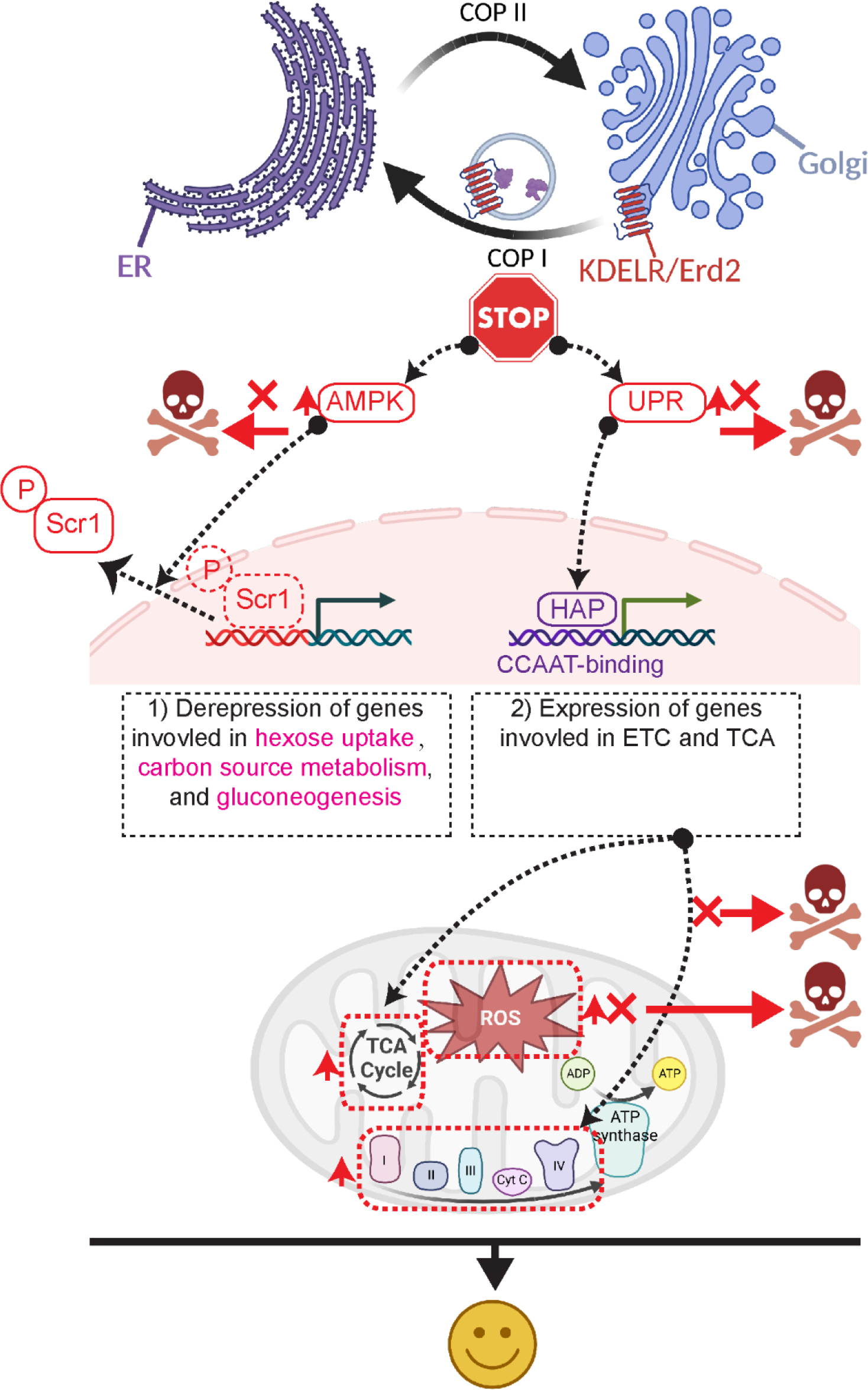
A model illustrating the alterations of the signaling pathways of UPR and AMPK and the responses of and mitochondria by the absence of KDELR/Erd2. The absence of Erd2 impairs the retrograde transport of the xDEL-containing ER resident proteins, leading to UPR and activation of AMPK. As a result, mitochondrial respiration and the production of mitochondrial ROS are enhanced by a transcriptional program that is controlled by UPR. In addition, the activation of AMPK derepress the genes involved in hexose uptake, carbon source metabolism and gluconeogenesis. Both UPR induction and AMPK activation are required for maintaining cell viability of cells lacking Erd2.

Erd2 collaborates with Ire1, the ER transmembrane sensor responsible for activating UPR, to maintain protein homeostasis in the ER of the budding yeast *Saccharomyces cerevisiae* (Beh and Rose, 1995). Yet whether the absence of Erd2 induces UPR had not been directly tested. In the present study, we provided evidence that UPR is induced by the absence of Erd2 (Fig. 2 A). The induction of UPR by the absence of Erd2 is likely due to the defective retention of the ER-stress response chaperones GRP78/Bip within the ER because GRP78/Bip is a typical cargo protein of KDELR/Erd2 (Capitani and Sallese, 2009; Dean and Pelham, 1990; Semenza et al., 1990) and functions to inhibit Ire1 in the ER (Preissler and Ron, 2019). In addition to GRP78/Bip, other cargo proteins of KDELR/Erd2, including protein disulfide isomerases, are required for promoting protein folding in the ER (Munro and Pelham, 1987). Therefore, it is also possible that the induction of UPR by the absence of Erd2 is due to the defective retention of the ER resident proteins required for promoting protein folding. In mammalian cells, expression of the KDELR mutant that fails to bind cargos enhances the responses of UPR induced with tunicamycin or dithiothreitol (Yamamoto et al., 2003). Hence, induction of UPR by malfunction of the KDELR/Erd2 retrieval system may be an evolutionarily conserved mechanism.

Our work also establishes that Erd2 regulates mitochondrial function in an UPR-dependent manner. Specifically, in cells lacking Erd2, the induction of UPR elevates the expression of the genes involved in the mitochondrial electron transport chain and the mitochondrial ATP synthase (Fig. 4 A and B and Fig. S2-S4). The elevation in gene expression is due to the increased expression of the transcriptional HAP complex. The reasons are as follows. It has been established that the transcriptional HAP complex, which binds to the CCAAT box within gene promoters (Mao and Chen, 2019), promotes expression of the genes involved in the mitochondrial electron transport and the tricarboxylic acid cycle (Buschlen et al., 2003; Takuma et al., 2013). Our findings revealed that the absence of Erd2 increased, in an Ire1/UPR-dependent manner, expression of the transcriptional HAP complex (Fig. 4 D). Moreover, the UPR inducer tunicamycin promotes expression of the transcriptional HAP complex (Fig. 4 A).

Of note, the absence of Erd2 induces UPR (Fig. 2 A). Therefore, our results support the model in which Erd2 alters mitochondrial function by guiding the expression of the UPR-promoted genes involved in the mitochondrial electron transport chain and the mitochondrial ATP synthase (Fig. 8).

Consistent with the finding that the absence of Erd2 increased expression of genes involved in mitochondrial electron transport chain and the mitochondrial ATP synthase, both the absence of Erd2 and the chemical induction of UPR by tunicamycin increased mitochondrial respiration and mitochondrial ROS (Fig. 1 B-D and Fig. 2 D and E). Therefore, these results further support the model in which activation of UPR boosts mitochondrial function and Erd2 regulates mitochondrial function via UPR. The model is consistent with two findings made with mammalian cells. One finding showed that the PERK arm of UPR promotes formation of mitochondrial hyperfusion to protect mitochondria under ER stresses (Lebeau et al., 2018), while the other finding showed that the PERK arm of UPR promotes formation of the mitochondrial respiratory complexes (Balsa et al., 2019). Hence, KDELR/Erd2 regulates mitochondrial function via UPR may be a mechanism that has been conserved through evolution.

In addition to induction of UPR, impairment of the KDELR/Erd2 retrieval system activates AMPK. This conclusion is supported by evidence that AMPK is phosphorylated in cells lacking Erd2 (Fig. 5 C and D). Moreover, the transcription factor Scr1, which functions to repress hexose uptake, carbon metabolism, and gluconeogenesis (Vassiliadis et al., 2019), is exported from the nucleus to the cytoplasm in cells lacking Erd2 (Fig. 5 A and B), a phenotype resulted from activated AMPK under glucose-limiting conditions (Matsuzawa et al., 2012). AMPK functions not only as a sensor for monitoring energy metabolism but also as a glucose sensor (Steinberg and Hardie, 2023). In general, under low energy or glucose-limiting conditions, AMPK is activated to promote catabolism but to inhibit anabolism (Steinberg and Hardie, 2023). Therefore, the activation of AMPK in cells lacking Erd2 indicates that *erd2*-deleted cells are at a low energy/glucose state. We noticed that the phosphorylation levels of AMPK in cells lacking Erd2 were significantly lower than that in cells cultured under glucose-starved conditions (Fig. 5 C and D). This result indicates that the absence of Erd2 may only partially activate AMPK, and such partial activation of AMPK is beneficial for maintaining cell viability of cells lacking Erd2 (Fig. 7).

The UPR-induced increases in mitochondrial respiration and ROS in cells lacking Erd2 are required for maintaining cell viability (Fig. 6). Given the role of the KDELR/Erd2 retrieval system in maintaining protein homeostasis in the ER, we speculated that the absence of Erd2 may increase energy demands required for restoring protein homeostasis. To meet the energy demands, mitochondria functions may be enhanced. This speculation predicts that defects in promoting mitochondrial functions are detrimental for the cells losing the KDELR/Erd2 retrieval system. Indeed, clearance of mitochondrial ROS (Fig. 6 B and D) or inhibition of mitochondrial respiration (Fig. 6 C and E) in cells lacking Erd2 caused cell death. Since the absence of Erd2 increases mitochondrial respiration and ROS in a UPR-dependent manner. The above speculation also predicts that defects in UPR should impair cell growth. Consistently, under unstressed conditions, *erd2*Δ*ire1*Δ cells, in which both mitochondrial respiration and mitochondrial ROS were diminished (Fig. 3 B-E), grew poorly but WT and *erd2*Δ grew comparably (Fig. 2 B).

In conclusion, we revealed a previously uncharacterized role of KDELR/Erd2 in guiding the signaling pathways of UPR and AMPK to maintain cellular homeostasis. Since KDELR/Erd2 have pathophysiological significance, our work provides insights into developing strategies for intervening in physiological disorders caused by malfunction of KDELR/Erd2.

## Materials and methods

### Yeast strains and plasmids

Yeast strains were created either by random spore digestion or tetrad-dissection. To create strains carrying deletion of a gene, the PCR-based homologous recombination method using the pFA6a series of plasmids was employed. Yeast strains used in this study are listed in supplementary Table S1. Plasmids were created by digestion with restriction enzymes purchased from New England Biolabs (www.neb.com) and ligation with T4 ligase (www.neb.com). Plasmids used in this study are listed in supplementary Table S2.

### Yeast cell culture

Yeast cells were cultured in Edinburgh Minimal Media supplemented with 2% glucose and five amino acids (adenine, leucine, uracil, histidine, and lysine, 0.225 g/L each) (referred to as EMM) at 30°C, and cells at the exponential phase were collected for analysis. All culture media were purchased from Formedium (www.formedium.com).

### Measurement of the rate of oxygen consumption

The rate of oxygen consumption was measured with a Strathkelvin respirometer (Model 782) (www.strathkelvin.com), according to the manufacturer’s instructions. First, yeast cells were cultured in EMM until the exponential phase (OD600=0.6-0.8) was reached. The measurements were then made at room temperature. Antimycin A (www.enzolifesciences.com) at a working concentration of 0.15 µg/ml was used to block mitochondrial respiration, which was performed in parallel and served as a control. Three independent experiments were performed.

### Measurement of mitochondrial reactive oxygen species

To visualize and quantify mitochondrial reactive oxygen species (mitochondrial ROS) of fission yeast cells, MitoSOX Red (www.invitrogen.com) was used at a working concentration of 5 µM. First, yeast cells were cultured in EMM until the exponential phase (OD600=0.6-0.8) was reached; MitoSOX was then added to the culture, followed by culture in the dark at 30°C for 30 minutes. In the case where ER stress was induced, tunicamycin (www.casmart.com.cn) at a working concentration of 0.5 µg/mL was added, and the tunicamycin-treated cells were cultured for 1 hour. To measure mitochondrial ROS of these cells, MitoSOX was added to the culture at 30 minutes before the end of tunicamycin treatment. Finally, yeast cells were collected and washed with fresh EMM, followed by observation with a spinning-disk microscope. Three independent experiments were performed.

To visualize and quantify mitochondrial ROS of HeLa cells, MitoSOX in Opti-MEM was added to the culture dish after cells were washed with PBS buffer three times; cells were then cultured at 37°C for 30 minutes. To stain DNA, Hoechest at a working concentration of 1 μg/ml was added to the culture dishes at 5 minutes before the end of MitoSOX staining. Once all staining procedures were completed, the original medium was removed and replaced with fresh DMEM medium, and cells were cultured at 37°C for 5 minutes, followed by microscopic observation. Three independent experiments were performed.

### Serial dilution growth assays

Yeast cells were cultured in liquid EMM medium until the exponential phase (OD600=0.6- 0.8) was reached. Serial dilutions (10-fold) of yeast cells were made, and then diluted cells were spotted onto solid EMM plates or EMM plates containing 0.5 µg/ml tunicamycin. The plates were incubated at 30 °C for three days. Three independent repeats were performed.

### Cell viability test

Cell viability was determined by tetrad-analysis of single cells. Specifically, yeast cells were cultured in EMM until the exponential phase (OD600=0.6-0.8) was reached. Before the drug treatment was conducted, yeast cells were collected, and single cells were separated and aligned on EMM plates (i.e., experiment at 0 hour). To clear ROS, Trolox at a working concentration of 1 mM was used to treat yeast cells for 4 or 8 hours at 30°C. To inhibit mitochondrial respiration, antimycin A at a working concentration of 0.15 µg/ml was used to treat yeast cells for 4 or 8 hours at 30°C. After the treatments, single yeast cells were separated and grown on EMM plates by tetrad-analysis. All plates were then incubated at 30°C for three days. Five independent experiments were performed.

### RNA extraction and real-time quantitative PCR

Yeast cells (1-5×10^7^) from 1-1.5 ml cultures were collected, and total RNA was extracted with a genomic DNA purification kit (R1002; www.zymoresearch.com). RT-PCR with a kit (HiScript III RT SuperMix, H6201080; www.yeasen.com) was then conducted to generate cDNAs.

Real-time quantitative PCR (qPCR) was accomplished with the primers listed in supplementary Table S3, using a kit purchased from YEASEN (Vazyme Univer Blue qPCR SYBR Master Mix, www.yeasen.com) on a Roche instrument (LightCycle 96, www.lifescience.roche.com). Five independent repeats were performed.

### RNA sequencing and analysis

RNA was extracted as described above. The concentration of RNA was determined by the NanoDrop (www.thermofisher.com) procedure. Three replicates were prepared for each type of strains. A total amount of 2 µg RNA per sample was used, and RNA sequencing was accomplished by Novogene (Beijing, China). Extracted RNAs were fragmented and then used for generating cDNAs with random primers. Subsequently, PCR was performed to amplify cDNAs with adaptor primers. An Illumina HiSeq instrument was used for sequencing.

Raw RNA-seq data were first processed by Trimmomatic (version 0.39) (Bolger et al., 2014) to trim off adaptors and eliminate reads that were in low quality. The cleaned RNA-seq data were aligned to the *Schizosaccharomyces pombe* reference genome GCF_000002945.1_ASM294v2.fa (Wood et al., 2012) by HISAT2 (version 2.2.0) (Kim et al., 2015). Gene expression levels were calculated by use of HTSeq (Anders et al., 2014), with gene models obtained from the reference file (GCF_000002945.1_ASM294v2_genomic.gff) derived from NCBI. Afterwards, analysis of differential gene expression was performed using a DESeq2 (version 1.26.0) package in R (See supplementary excel file). GO enrichment analysis on the differentially expressed genes identified by DESeq2 were conducted using Annotation Hub package (version 1.26.0) in R. In RNA sequencing, three independent replicates were performed for each type of strains.

### Microscopy and data analysis

Live-cell imaging was accomplished using a PerkinElmer Ultraview spinning disk microscope equipped with a Nikon Apochromat TIRF 100X 1.49NA objective and a Hamamatsu C9100-23B EMCCD camera, and 405-, 488-, and 561-nm lasers were used in the experiments (www.perkinelmer.com). Images were acquired by Volocity (www.perkinelmer.com). To observe yeast cells, stack images containing 11 planes at 0.5 μm spacing were acquired. Yeast cells were observed on slides coated with EMM agarose (3%) pads.

Microscopic data were analyzed with Fiji Image J 1.52 (www.imagej.nih.gov) and MetaMorph 7.7 (www.moleculardevices.com). Graphs were created by KaleidaGraph 4.5 (www.synergy.com). Normality of data was determined by OriginPro (version 2021b) (www.originlab.com), and statistical analysis was performed by KaleidaGraph 4.5. The illustration graph in Fig. 8 was created using graphs generated with BioRender.com.

## Acknowledgments

We thank members in the Fu laboratory for insightful discussion. This work is supported by grants from National Key Research and Development Program of China [2022YFA1303100], National Natural Science Foundation of China [91754106, 32070707, and 31621002], the Fundamental Research Funds for the Central Universities [WK9110000151], and the Center for Advanced Interdisciplinary Science and Biomedicine of IHM [QYPY20220003].

## Author contributions

C.F. conceived the project. M.Z., Z.F., Y.W., F.D., Y.W., F.Z., and J.H. performed experiments. M.Z. and Z.F. analyzed data. S.M., X.M., X.L., X.Y., and C.F. supervised the project. M.Z., D.H., X.Y., and C.F. wrote the manuscript.

## References

1. Anders, S., P.T. Pyl, and W. Huber. 2014. HTSeq—a Python framework to work with high-throughput sequencing data. Bioinformatics. 31:166–169.

2. Aoe, T., E. Cukierman, A. Lee, D. Cassel, P.J. Peters, and V.W. Hsu. 1997. The KDEL receptor, ERD2, regulates intracellular traffic by recruiting a GTPase-activating protein for ARF1. EMBO J. 16:7305-7316.

3. Balsa, E., M.S. Soustek, A. Thomas, S. Cogliati, C. Garcia-Poyatos, E. Martin-Garcia, M. Jedrychowski, S.P. Gygi, J.A. Enriquez, and P. Puigserver. 2019. ER and Nutrient Stress Promote Assembly of Respiratory Chain Supercomplexes through the PERK- eIF2alpha Axis. Mol Cell. 74:877–890 e876.

4. Beh, C.T., and M.D. Rose. 1995. Two redundant systems maintain levels of resident proteins within the yeast endoplasmic reticulum. Proc Natl Acad Sci U S A. 92:9820–9823.

5. Bolger, A.M., M. Lohse, and B. Usadel. 2014. Trimmomatic: a flexible trimmer for Illumina sequence data. Bioinformatics. 30:2114–2120.

6. Brauer, P., J.L. Parker, A. Gerondopoulos, I. Zimmermann, M.A. Seeger, F.A. Barr, and S. Newstead. 2019. Structural basis for pH-dependent retrieval of ER proteins from the Golgi by the KDEL receptor. Science. 363:1103–1107.

7. Buschlen, S., J.M. Amillet, B. Guiard, A. Fournier, C. Marcireau, and M. Bolotin-Fukuhara. 2003. The S. Cerevisiae HAP complex, a key regulator of mitochondrial function, coordinates nuclear and mitochondrial gene expression. Comp Funct Genomics. 4:37–46.

8. Cabrera, M., M. Muniz, J. Hidalgo, L. Vega, M.E. Martin, and A. Velasco. 2003. The retrieval function of the KDEL receptor requires PKA phosphorylation of its C-terminus. Mol Biol Cell. 14:4114–4125.

9. Calvo, I.A., N. Gabrielli, I. Iglesias-Baena, S. Garcia-Santamarina, K.L. Hoe, D.U. Kim, M. Sanso, A. Zuin, P. Perez, J. Ayte, and E. Hidalgo. 2009. Genome-wide screen of genes required for caffeine tolerance in fission yeast. PLoS One. 4:e6619.

10. Cancino, J., A. Capalbo, A. Di Campli, M. Giannotta, R. Rizzo, J.E. Jung, R. Di Martino, M. Persico, P. Heinklein, M. Sallese, and A. Luini. 2014. Control systems of membrane transport at the interface between the endoplasmic reticulum and the Golgi. Dev Cell. 30:280–294.

11. Capitani, M., and M. Sallese. 2009. The KDEL receptor: new functions for an old protein. FEBS Lett. 583:3863–3871.

12. Chiron, S., M. Gaisne, E. Guillou, P. Belenguer, G.D. Clark-Walker, and N. Bonnefoy. 2007. Studying mitochondria in an attractive model: Schizosaccharomyces pombe. Methods Mol Biol. 372:91–105.

13. Dean, N., and H.R. Pelham. 1990. Recycling of proteins from the Golgi compartment to the ER in yeast. J Cell Biol. 111:369–377.

14. Dong, F., M. Zhu, F. Zheng, and C. Fu. 2022. Mitochondrial fusion and fission are required for proper mitochondrial function and cell proliferation in fission yeast. FEBS J. 289:262–278.

15. Gerondopoulos, A., P. Brauer, T. Sobajima, Z. Wu, J.L. Parker, P.C. Biggin, F.A. Barr, and S. Newstead. 2021. A signal capture and proofreading mechanism for the KDEL-receptor explains selectivity and dynamic range in ER retrieval. Elife. 10.

16. Guydosh, N.R., P. Kimmig, P. Walter, and R. Green. 2017. Regulated Ire1-dependent mRNA decay requires no-go mRNA degradation to maintain endoplasmic reticulum homeostasis in S. pombe. Elife. 6.

17. Hamada, H., M. Suzuki, S. Yuasa, N. Mimura, N. Shinozuka, Y. Takada, M. Suzuki, T. Nishino, H. Nakaya, H. Koseki, and T. Aoe. 2004. Dilated cardiomyopathy caused by aberrant endoplasmic reticulum quality control in mutant KDEL receptor transgenic mice. Mol Cell Biol. 24:8007–8017.

18. Hardwick, K.G., J.C. Boothroyd, A.D. Rudner, and H.R. Pelham. 1992. Genes that allow yeast cells to grow in the absence of the HDEL receptor. EMBO J. 11:4187–4195.

19. Hardwick, K.G., M.J. Lewis, J. Semenza, N. Dean, and H.R. Pelham. 1990. ERD1, a yeast gene required for the retention of luminal endoplasmic reticulum proteins, affects glycoprotein processing in the Golgi apparatus. EMBO J. 9:623–630.

20. Kamimura, D., K. Katsunuma, Y. Arima, T. Atsumi, J.J. Jiang, H. Bando, J. Meng, L. Sabharwal, A. Stofkova, N. Nishikawa, H. Suzuki, H. Ogura, N. Ueda, M. Tsuruoka, M. Harada, J. Kobayashi, T. Hasegawa, H. Yoshida, H. Koseki, I. Miura, S. Wakana, K. Nishida, H. Kitamura, T. Fukada, T. Hirano, and M. Murakami. 2015. KDEL receptor 1 regulates T-cell homeostasis via PP1 that is a key phosphatase for ISR. Nat Commun. 6:7474.

21. Kim, D., B. Langmead, and S.L. Salzberg. 2015. HISAT: a fast spliced aligner with low memory requirements. Nat Methods. 12:357–360.

22. Kimmig, P., M. Diaz, J. Zheng, C.C. Williams, A. Lang, T. Aragon, H. Li, and P. Walter. 2012. The unfolded protein response in fission yeast modulates stability of select mRNAs to maintain protein homeostasis. Elife. 1:e00048.

23. Lebeau, J., J.M. Saunders, V.W.R. Moraes, A. Madhavan, N. Madrazo, M.C. Anthony, and R.L. Wiseman. 2018. The PERK Arm of the Unfolded Protein Response Regulates Mitochondrial Morphology during Acute Endoplasmic Reticulum Stress. Cell Rep. 22:2827–2836.

24. Lemos, C., V.K. Schulze, S.J. Baumgart, E. Nevedomskaya, T. Heinrich, J. Lefranc, B. Bader, C.D. Christ, H. Briem, L.P. Kuhnke, S.J. Holton, U. Bomer, P. Lienau, F. von Nussbaum, C.F. Nising, M. Bauser, A. Hagebarth, D. Mumberg, and B. Haendler. 2021. The potent AMPK inhibitor BAY-3827 shows strong efficacy in androgen-dependent prostate cancer models. Cell Oncol (Dordr). 44:581–594.

25. Lewis, M.J., and H.R. Pelham. 1990. A human homologue of the yeast HDEL receptor. Nature. 348:162–163.

26. Lewis, M.J., and H.R. Pelham. 1992. Ligand-induced redistribution of a human KDEL receptor from the Golgi complex to the endoplasmic reticulum. Cell. 68:353–364.

27. Lewis, M.J., D.J. Sweet, and H.R. Pelham. 1990. The ERD2 gene determines the specificity of the luminal ER protein retention system. Cell. 61:1359–1363.

28. Ling, N.X.Y., A. Kaczmarek, A. Hoque, E. Davie, K.R.W. Ngoei, K.R. Morrison, W.J. Smiles, G.M. Forte, T. Wang, S. Lie, T.A. Dite, C.G. Langendorf, J.W. Scott, J.S. Oakhill, and J. Petersen. 2020. mTORC1 directly inhibits AMPK to promote cell proliferation under nutrient stress. Nat Metab. 2:41–49.

29. Mao, Y., and C. Chen. 2019. The Hap Complex in Yeasts: Structure, Assembly Mode, and Gene Regulation. Front Microbiol. 10:1645.

30. Matsuzawa, T., Y. Fujita, H. Tohda, and K. Takegawa. 2012. Snf1-like protein kinase Ssp2 regulates glucose derepression in Schizosaccharomyces pombe. Eukaryot Cell. 11:159–167.

31. Munro, S., and H.R. Pelham. 1987. A C-terminal signal prevents secretion of luminal ER proteins. Cell. 48:899–907.

32. Preissler, S., and D. Ron. 2019. Early Events in the Endoplasmic Reticulum Unfolded Protein Response. Cold Spring Harb Perspect Biol. 11.

33. Rasul, F., F. Zheng, F. Dong, J. He, L. Liu, W. Liu, J.Y. Cheema, W. Wei, and C. Fu. 2021. Emr1 regulates the number of foci of the endoplasmic reticulum-mitochondria encounter structure complex. Nat Commun. 12:521.

34. Robinson, D.G., and F. Aniento. 2020. A Model for ERD2 Function in Higher Plants. Front Plant Sci. 11:343.

35. Scheel, A.A., and H.R. Pelham. 1998. Identification of amino acids in the binding pocket of the human KDEL receptor. J Biol Chem. 273:2467–2472.

36. Semenza, J.C., K.G. Hardwick, N. Dean, and H.R. Pelham. 1990. ERD2, a yeast gene required for the receptor-mediated retrieval of luminal ER proteins from the secretory pathway. Cell. 61:1349–1357.

37. Semenza, J.C., and H.R. Pelham. 1992. Changing the specificity of the sorting receptor for luminal endoplasmic reticulum proteins. J Mol Biol. 224:1–5.

38. Siggs, O.M., D.L. Popkin, P. Krebs, X. Li, M. Tang, X. Zhan, M. Zeng, P. Lin, Y. Xia, M.B. Oldstone, R.J. Cornall, and B. Beutler. 2015. Mutation of the ER retention receptor KDELR1 leads to cell-intrinsic lymphopenia and a failure to control chronic viral infection. Proc Natl Acad Sci U S A. 112:E5706–5714.

39. Steinberg, G.R., and D.G. Hardie. 2023. New insights into activation and function of the AMPK. Nat Rev Mol Cell Biol. 24:255–272.

40. Takuma, K., H. Ohtsuka, K. Azuma, H. Murakami, and H. Aiba. 2013. The fission yeast php2 mutant displays a lengthened chronological lifespan. Biosci Biotechnol Biochem. 77:1548–1555.

41. Townsley, F.M., G. Frigerio, and H.R. Pelham. 1994. Retrieval of HDEL proteins is required for growth of yeast cells. J Cell Biol. 127:21–28.

42. Townsley, F.M., D.W. Wilson, and H.R. Pelham. 1993. Mutational analysis of the human KDEL receptor: distinct structural requirements for Golgi retention, ligand binding and retrograde transport. EMBO J. 12:2821–2829.

43. Trychta, K.A., S. Back, M.J. Henderson, and B.K. Harvey. 2018. KDEL Receptors Are Differentially Regulated to Maintain the ER Proteome under Calcium Deficiency. Cell Rep. 25:1829–1840 e1826.

44. Vassiliadis, D., K.H. Wong, A. Andrianopoulos, and B.J. Monahan. 2019. A genome-wide analysis of carbon catabolite repression in Schizosaccharomyces pombe. BMC Genomics. 20:251.

45. Wilson, D.W., M.J. Lewis, and H.R. Pelham. 1993. pH-dependent binding of KDEL to its receptor in vitro. J Biol Chem. 268:7465–7468.

46. Wu, H., B.S. Ng, and G. Thibault. 2014. Endoplasmic reticulum stress response in yeast and humans. Biosci Rep. 34.

47. Yamamoto, K., H. Hamada, H. Shinkai, Y. Kohno, H. Koseki, and T. Aoe. 2003. The KDEL receptor modulates the endoplasmic reticulum stress response through mitogen-activated protein kinase signaling cascades. J Biol Chem. 278:34525–34532.

48. Zheng, F., B. Jia, F. Dong, L. Liu, F. Rasul, J. He, and C. Fu. 2019. Glucose starvation induces mitochondrial fragmentation depending on the dynamin GTPase Dnm1/Drp1 in fission yeast. J Biol Chem. 294:17725–17734.

