## Supplemental Figures and Tables for "A KDELR-mediated ER retrieval system guides the signaling pathways of UPR and AMPK to maintain cellular homeostasis"

**This PDF file includes:**

Supplementary Figures S1 to S4
Tables S1 to S3

### Supplementary figures and figure legends

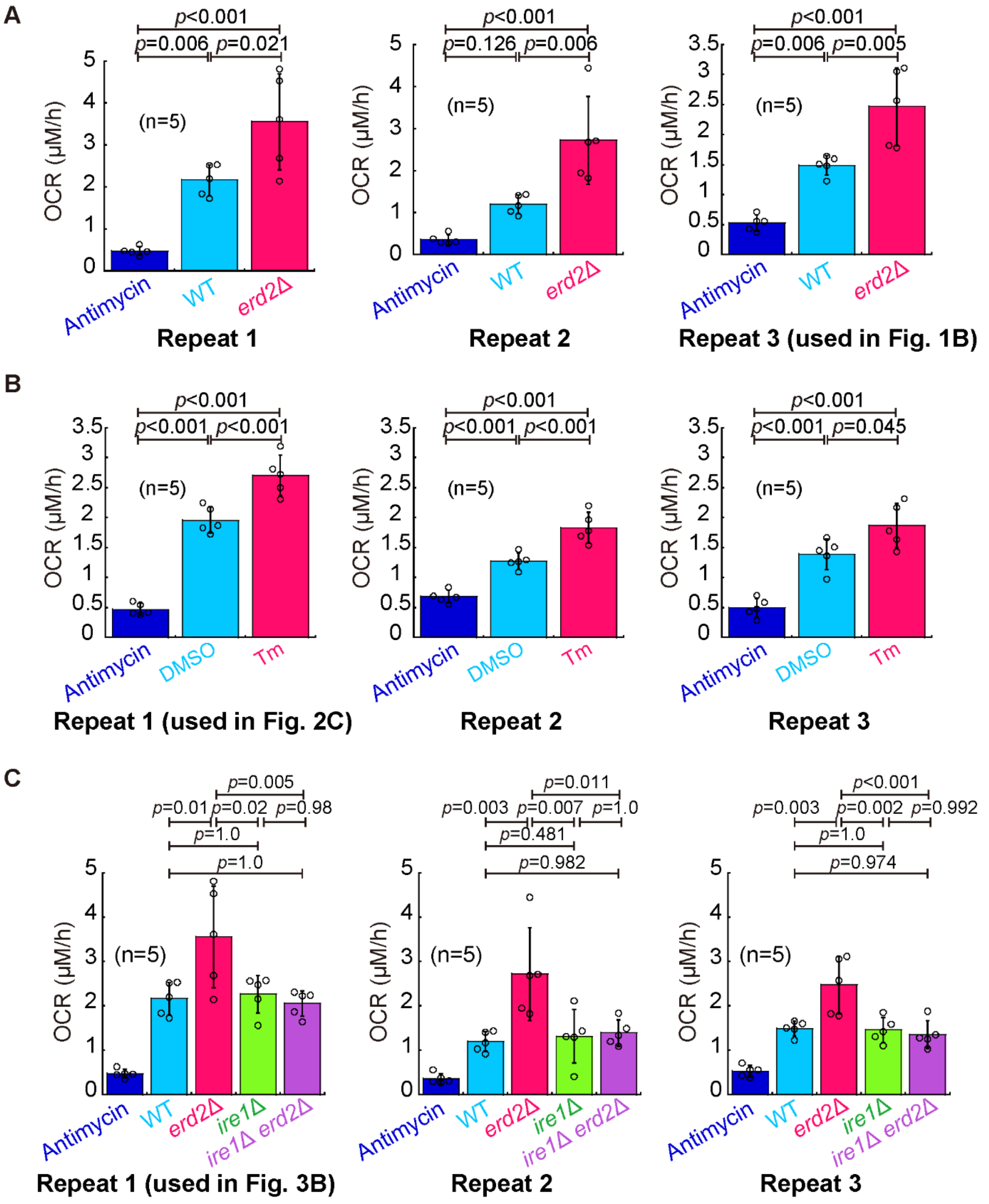


**Supplementary Fig. S1. Oxygen consumption rates (OCR) of the indicated cells (Related to main Fig. 1B, 2D, and 3B).** (A) A Strathkelvin oxygen respirometer was used to measure OCR. Shown are the results obtained from three independent experiments (repeat 3 was used in main Fig. 1B), and 5 repeats were performed for each type of cells. The height of the column is the mean, and error bars indicate S.D. The *p* values were calculated by one-way ANOVA with a Post Hoc Tukey HSD test.

Antimycin A (0.15 µg/ml) treatment was used inhibit mitochondrial respiration. (B) A Strathkelvin oxygen respirometer was used to measure OCR. Shown are the results obtained from three independent experiments (repeat 1 was used in main Fig. 2D), and 5 repeats were performed for each type of treatments. The height of the column is the mean, and error bars indicate S.D. The *p* values were calculated by one-way ANOVA with a Post Hoc Tukey HSD test. Antimycin A (0.15 µg/ml) treatment was used to inhibit mitochondrial respiration. (C) A Strathkelvin oxygen respirometer was used to measure OCR. Shown are the results obtained from three independent experiments (repeat 1 was used in main Fig. 3B), and 5 repeats were performed for each type of cells. The height of the column is the mean, and error bars indicate S.D. The *p* values were calculated by one-way ANOVA with a Post Hoc Tukey HSD test. Antimycin A (0.15 µg/ml) treatment was used to inhibit mitochondrial respiration.

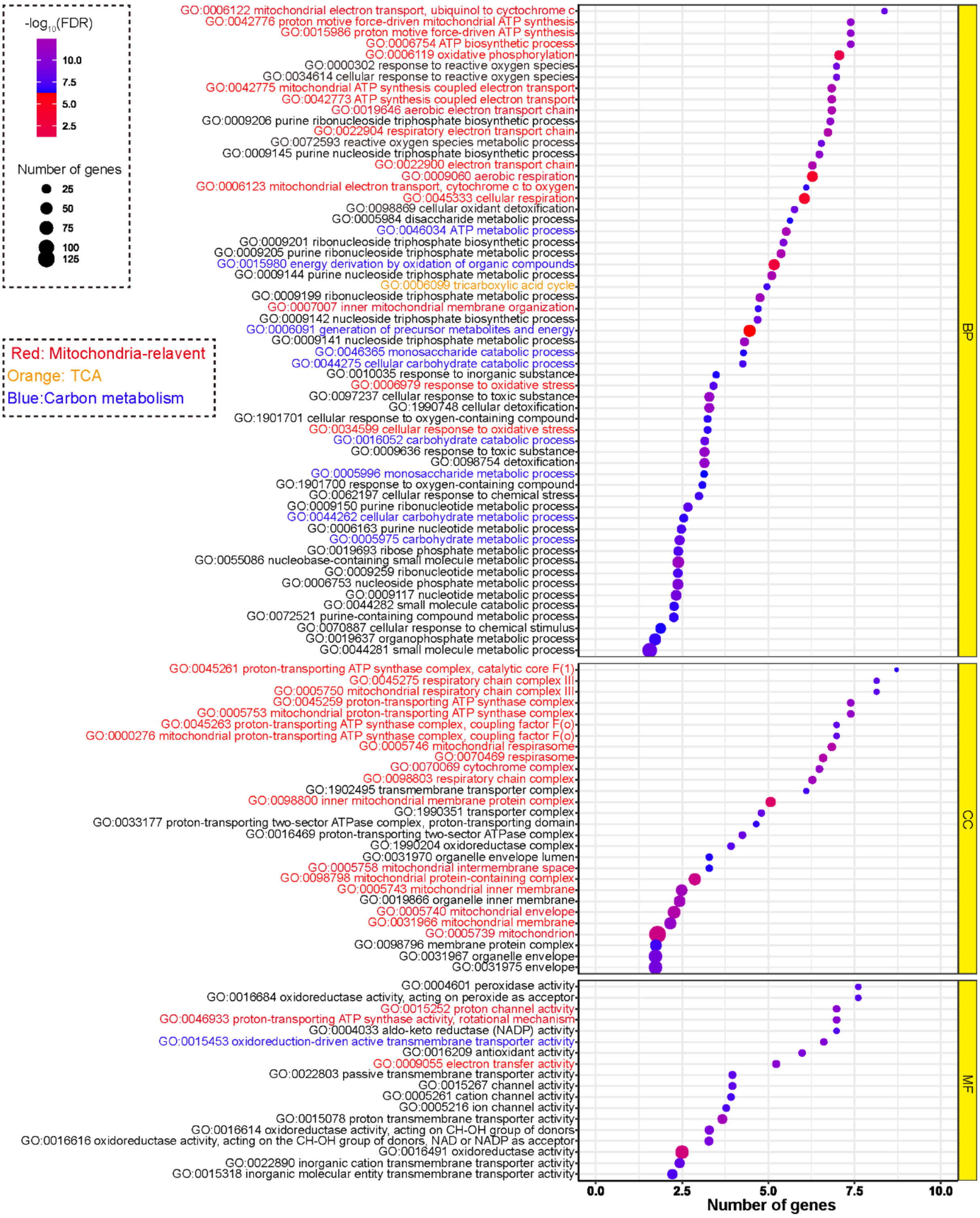


**Supplementary Fig. S2. GO enrichment analysis of the differentially expressed genes between *erd2*Δ and WT (Related to Fig. 4).** GO enrichment analysis. The differentially expressed genes (fold changes>1.5) identified by DESeq2 were analyzed by the Enrichr package with the *Schizosaccharomyces pombe* annotation file from Annotation Hub R package (version 1.26.0) and the results were presented by bubble plots. *MF*, molecular function; *BP*, biological process; *CC*, cellular component.

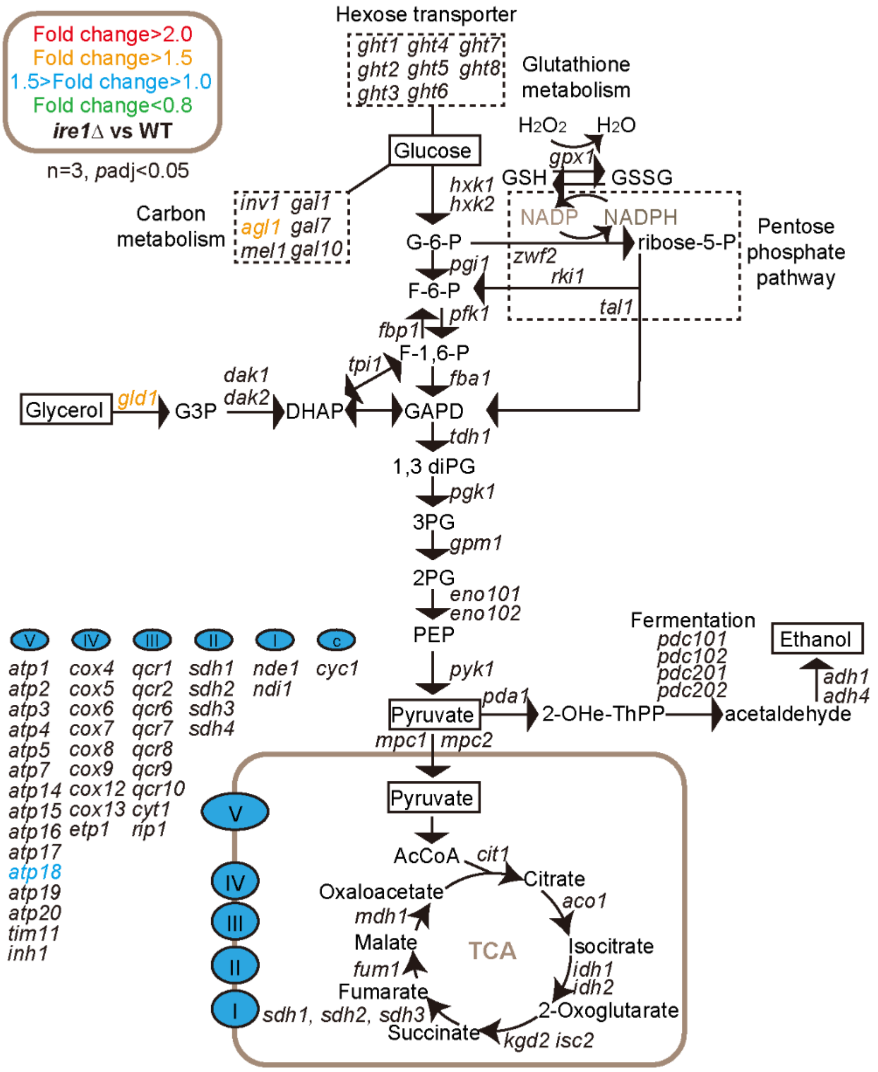


**Supplementary Fig. S3. Differentially expressed genes between *ire1*Δ and WT (Related to Fig. 4A).** Diagram illustrating glycolysis, the mitochondrial electron transport chain (ETC), and the tricarboxylic acid cycle (TCA). Indicated, in red, orange, blue, and green colors, are the genes that exhibit differential expression in *ire1*Δ cells (vs WT, adjusted p-values (padj)<0.05). The genes of the mitochondrial electron transport chain and the ATP synthetase are listed.

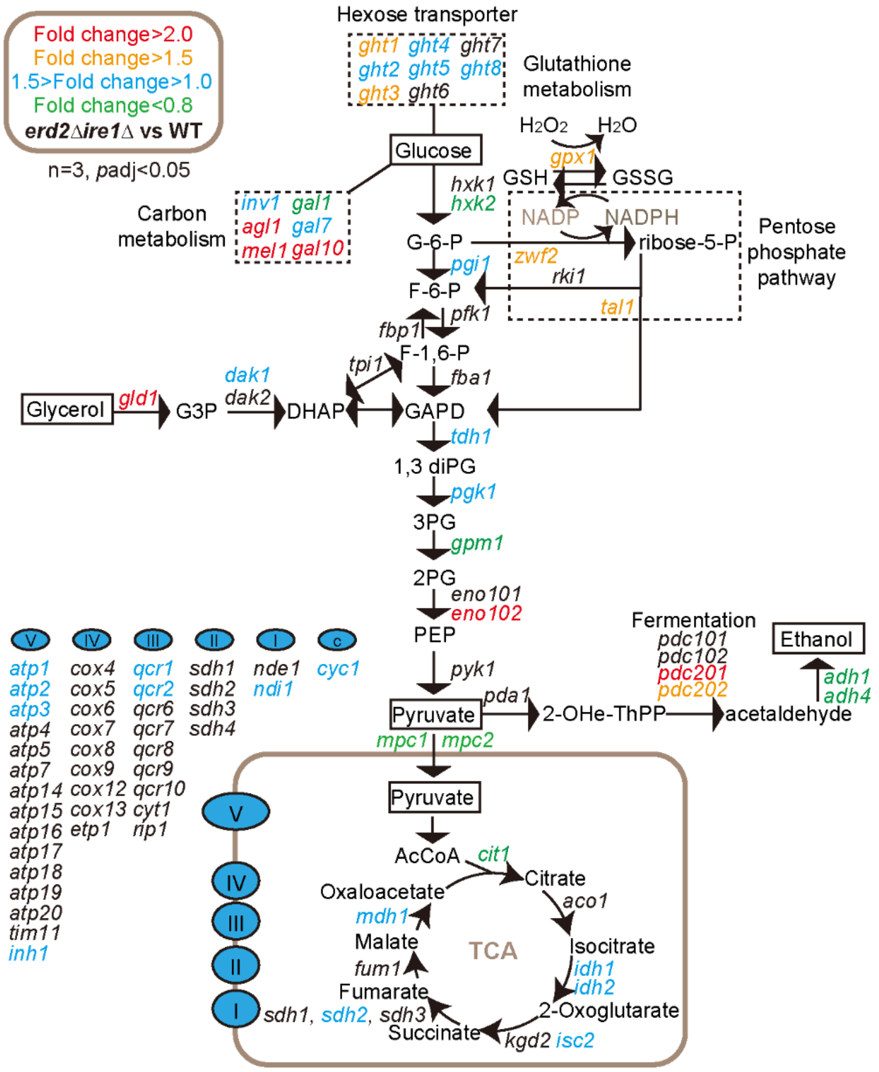


**Supplementary Fig. S4. Differentially expressed genes between *ire1*Δ*erd2*Δ and WT (Related to Fig. 4A).** Diagram illustrating glycolysis, the mitochondrial electron transport chain (ETC), and the tricarboxylic acid cycle (TCA). Indicated, in red, orange, blue, and green colors, are the genes that exhibit differential expression in *ire1*Δ*erd2*Δ cells (vs WT, adjusted p-values (padj)<0.05). The genes of the mitochondrial electron transport chain and the ATP synthetase are listed.

### Supplementary tables

#### Table S1. Yeast strains

| Strain | Genotype | Source |
|---|---|---|
| CF.4096 | WT h- | Laboratory collection |
| CF.11389 | *erd2*Δ:NatR h- | This study |
| CF.11390 | *ire1*Δ:HygR h- | This study |
| CF.11450 | *erd2*Δ:NatR *ire1*Δ:HygR h- | This study |

#### Table S2. Plasmids

| Plasmid | Genotype | Source |
|---|---|---|
| pCF.2511 | pEGFP | Laboratory collection |
| pCF.4536 | pEGFP-KDEL | This study |

#### Table S3. Primers used for the RT-PCR experiments

| Number | Name | Sequence |
|---|---|---|
| oCF.3355 | gas2-RT-F | TTACTTTGCAGCTGTTGTCGG |
| oCF.3356 | gas2-RT-R | CCTCGGCTAAGGGATCAACG |
| oCF.4327 | yop1-RT-F | ACAGACACTTGATCCGTCCC |
| oCF.4328 | yop1-RT-R | GCAAAAGAAGAGGCTGTGGG |
| oCF.4973 | Php2-RT-F | ATCCATATGAGCCGGTCGAG |
| oCF.4974 | Php2-RT-R | AGTCAAGAATCGGCCACCAG |
| oCF.4975 | Php3-RT-F | ACTGCGTCCAAGATTGCGTT |
| oCF.4976 | Php3-RT-R | TCTTCGGACCGGCTTTGTTTA |
| oCF.4977 | Php4-RT-F | CCGTGAATCCGAGGAAACCA |
| oCF.4978 | Php4-RT-R | AAGTGACGGGACGAACATCG |
| oCF.4979 | Php5-RT-F | ACAAGGCTTTCCCGAAGGTT |
| oCF.4980 | Php5-RT-R | ACCTCAATGGGTGCACTAGC |
| oCF.6357 | Atp1-RT-F | GGAACGTATTCGTGGAGCCT |
| oCF.6358 | Atp1-RT-R | ACAACCAACGGTATCGGCTT |
| oCF.6359 | Atp2-RT-F | GTAGATGAACGCGGTCCCAT |
| oCF.6360 | Atp2-RT-R | ACACCTGCACCACCGAATAG |
| oCF.6363 | cox13-RT-F | GAAGAGGAGCATGCAAAGCG |
| oCF.6364 | cox13-RT-R | TCTTGCTACCGTCTCCCCAT |
| oCF.6365 | rip1-RT-F | GCTTCCCGTTTCAAGCACTG |

| | | |
|---|---|---|
| oCF.6366 | rip1-RT-R | AGAGCACCCATTGTGCCTAC |
